# A model of protein folding with multiple native states: Metamorphicity, Intrinsic disorderness and folding upon binding of proteins

**DOI:** 10.1101/2025.02.26.640301

**Authors:** Rohon Mitra, Biman Jana

## Abstract

The sequence-structure-function paradigm in biology states that protein’s amino acid sequence determines its unique folded state structure which in turn dictates its unique biological function. This classic concept has been severely challenged by the discovery of metamorphic and intrinsically disorder proteins (IDPs). Metamorphic proteins can fold into multiple native structures and perform multiple functions. IDPs, on the contrary, remain unstructured at physiological conditions, but can fold to unique structure upon binding to a target protein and show functionality. Here, we present a statistical mechanics model of protein folding with multiple native states and analyze their folding phase diagrams. While recovering classic sequence-structure-function paradigm for single native state, our model shows metamorphicity at lower number of native states. An expansion of unfolded region in the phase diagram at higher number of native states, making unfolded state as the stable state at physiological conditions, indicates the emergence of IDP-like scenario. Folding upon binding scenario of IDPs has also been demonstrated when an energetic bias is introduced for a specific native state. A regaining of folded region upon biasing, increasing the folding propensity of the system, with folding towards a specific native state is shown. Therefore, our model is general enough to reproduce the classic sequence-structure-function, metamorphicity, intrinsic disorderness and folding upon binding scenarios of proteins.

**Significance Statement:** Proteins exhibit remarkable structural diversity, from having unique single native structure to having no particular structure but an inherently disordered ensemble that usually attains a single structure upon binding to another target. Here, we present a general statistical mechanics model of protein folding that reproduces single native state folding, metamorphicity, intrinsic disorder, and binding-induced folding by incorporating multiple native states. A notable increase in propensity of being in unfolded state, resembling intrinsic disorder at physiological conditions, for system with larger number of native state was shown. Introduction of a bias was shown to increase the folding propensity of such proteins directed toward a specific native state. The model’s ability to capture a wide range of structural diversity in proteins makes it highly versatile.

## Introduction

Protein folding is an intricate and highly regulated process that involves the transformation of a linear chain of amino acids into a specific, functional three-dimensional structure. The Anfinsen hypothesis [1] states that the final structure of a protein is determined by its amino acid sequence, meaning that the sequence contains all the information needed for it to fold properly under the right conditions [2, 3]. This process is essential for the proper functioning of proteins, as the final folded structure determines the ability of the protein to interact with other molecules and perform its biological functions [4–8]. On the issue of folding timescale, Levinthal’s paradox [9] points out that randomly sampling all possible conformations would take an impractically long time for a protein to reach its native state. However, despite the many possible conformations, a protein efficiently finds its correct structure within a biologically relevant timescale. Therefore, proteins do not fold through random sampling; rather, the amino acid sequence encodes the necessary information to guide the folding process along specific energetically favorable pathways [10–12], navigate through a complex energy landscape [13–15] that directs the protein towards its stable native state by avoiding high-energy barriers [16,17] and favoring low-energy conformations. The sequence dictates how the protein will fold, and certain regions form secondary structures such as alpha helices and beta sheets, which then assemble in the final tertiary structure [18]. In recent years, advances in computational approaches, particularly in sequence-to-structure prediction, have made significant progress toward understanding protein folding. One of the most notable breakthroughs is AlphaFold, an AI system that predicts protein structures from amino acid sequences with unprecedented precision [19, 20]. However, while AlphaFold successfully predicts native structures, it does not reveal the folding pathway [21, 22], the step-by-step process by which the protein achieves its functional form. When the folding process goes wrong, proteins can misfold [23–25] and aggregate [26], leading to dysfunctional proteins associated with diseases [27], such as Alzheimer’s [28] and Parkinson’s [29, 30].

Most recently, the concept of one sequence providing one unique structure/function is challenged. Certain proteins, such as metamorphic proteins [31–33], can adopt multiple stable conformations, allowing them to switch between functional states depending on environmental conditions [34,35]. Another important class of proteins, Intrinsically disordered proteins (IDPs) do not even fold into fixed three-dimensional structures but remain flexible, exist in various conformations [36,37], and often play a critical role in signaling and regulation. However, these IDPs can adopt specific three-dimensional structures when bound to specific target proteins and can perform multiple functions [38, 39].

To better understand protein folding, various physics-based models have been developed, each capturing different aspects of the folding process. One such foundational model is the spin-glass framework [40–45], originally developed for disordered magnetic systems. In their seminal work, Bryngelson and Wolynes have developed a theoretical model of protein folding with a single native state in a modified spin-glass framework. [46] Using a mean-field approach with random energy approximation, the model provides a phase diagram for protein with 3 main phases: the unfolded disordered state, the native folded state, and a glassy phase. [47, 48] The model reveals that, under certain conditions, the system undergoes a first-order phase transition between folded and unfolded states. Furthermore, the glassy phase suggests the presence of a second-order phase transition [49] in the system as well. It captures the complexity of rugged energy landscapes with numerous local minima. However, a theoretical framework that includes scenarios like metamorphic proteins, IDPs, and folding upon binding of IDPs is still lacking.

In this work, we present a theoretical model to investigate the folding behavior of proteins with multiple native states. We first develop a generalized framework for systems with *n* native states and analyze the phase diagram features for *n* = 1 to 5. The results are correlated with metamorphic proteins and intrinsically disordered proteins (IDPs). Moreover, our model naturally captures folding-upon-binding scenarios observed in IDPs.

## Model

We base our approach on the spin-glass model of protein folding, extending it to systems with multiple native states, as first proposed by Wolynes and co-workers [46, 50] in their seminal work on protein folding dynamics. In this model, the folding process is governed by a complex energy landscape with multiple local minima, where proteins navigate through various configurations to reach their folded functional state. To describe the total energy of a configuration of a protein, different energy terms were defined:*ε*_*i*_(*α*_*i*_) represents the energy of *i*^th^ amino acid residue in its specific conformation, *α*_*i*_. *J*_*i*,*i*+1_(*α*_*i*_, *α*_*i*+1_) accounts for nearest-neighbor interactions between two successive amino acids in the peptide chain, and *K*_*i*,*j*_(*α*_*i*_, *α*_*j*_, *r*_*i*_, *r*_*j*_) describes long-range interactions between residues that are spatially close in the tertiary structure of that configuration of the protein. Here, *α*_*i*_ denotes the state of a particular residue at position *r*_*i*_ in the sequence, and each residue can adopt one native state along with approximately 10 non-native states(*ν* ≈ 10), as originally suggested by Wolynes and co-workers.

The total energy of the protein in a particular configuration is therefore expressed as:

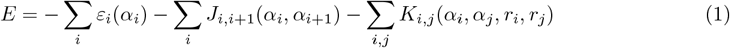

However, working with Eq. (1) analytically is highly challenging due to the complexity introduced by a large number of variables. A stochastic Hamiltonian approach is introduced to statistically model energy states, capturing the essential features of spin-glass behavior. In line with Derrida’s random energy approximation, they assumed that the energies of different protein conformations are uncorrelated, simplifying the model while maintaining its key characteristics. Considering a protein with *N* residues, where *N*_0_ number of residues are in their native state, there can be different possible protein configurations with different total energies. The joint probability distribution for *n* configurations with energies *E*_1_, *E*_2_, …, *E*_*n*_ is given by:

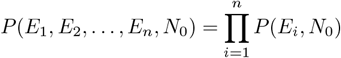

At the residue level, non-native configuration energies are considered to be with mean 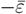 and standard deviation of Δ*ε*, while all native configuration energies are set at −*ε*_0_, where 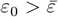. For the nearestneighbor interaction term, non-native interactions have a mean of 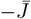 and standard deviation of Δ*J*, with all native interactions set to −*J*, where 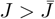. At the tertiary level, non-native interactions follow a distribution with mean 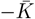 and standard deviation of Δ*K*, while all native interactions are set to −*K*, where 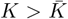. Typically, the number of tertiary interactions for each residue (z) is around 2–3 for protein, with *z* varying slightly between the completely unfolded and fully folded conformations. The random energy approximation applies to these structural interactions and the distribution of total energy with *N*_0_ number of residues are in their native state is modeled as a Gaussian distribution with the average being:

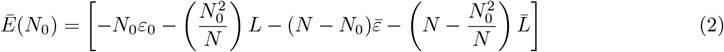

where:

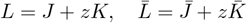

The standard deviation of the distributionΔ*E*(*N*_0_) is given by:

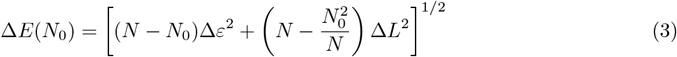

where:

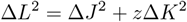

In our model, while keeping the basic parameters unchanged, we assume that each amino acid residue can exist in more than one native state and ten non-native states. Let us assume a situation where *N*_0_ number of residues are in the native state out of total N residues. The total count of native residues will now be distributed across *n* alternative native states, with *N*_*i*_ residues in the *i*-th native state. The combined total of residues across all native states satisfies the equation:

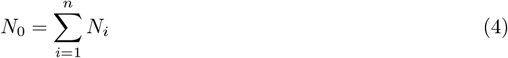

.The energies of different protein configuration under these condition is a Gaussian probability distribution *P* (*E, N*_1_, *N*_2_, …, *N*_*n*_), where *N*_*i*_ represents the number of residues in the *i*-th distinct native state. The average energy of this distribution is given by

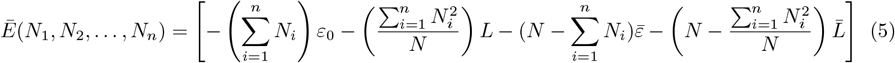

The standard deviation is expressed as

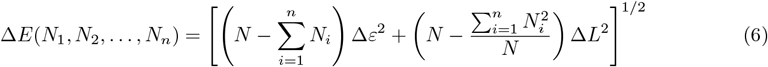

By comparing Eqs. 2 and 3 with Eqs. 5 and 6, it becomes evident that, for a given value of *N*_0_, the multi-state model exhibits a lower average value of the distribution compared to the single-state model. Additionally, the standard deviation is larger in the multi-state model, reflecting greater variability. The density of states, denoted by *n*(*E*), is a critical parameter in our study. We sum the density of states over all possible values of *N*_*i*_ for *i* = 1, …, *n* to obtain the average density of states ⟨*n*(*E*)⟩ for the multi-state system, expressed as follows:

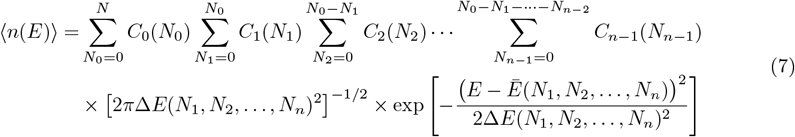

where

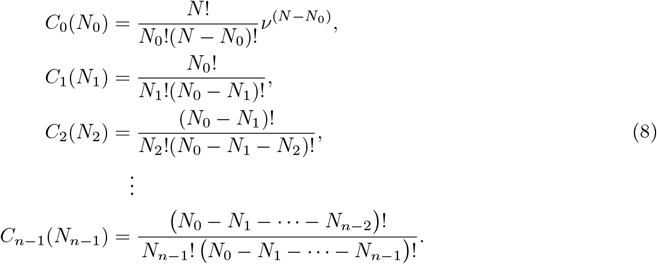

The coefficient *C*_0_(*N*_0_) represents the combinatorial expression for selecting *N*_0_ random native residues from the total of *N* residues in the model protein system. The coefficient *C*_*i*_(*N*_*i*_) represents the combinatorial factor for selecting *N*_*i*_ specific residues from the total *N*_0_ native-like residues for each *i* = − 1, 2, …, *n* 1. The sum of all values of *n*(*E*) will receive significant contributions from various terms. However, in the thermodynamic limit, when *N* is large, the overall behavior of the model system is predominantly governed by the largest term in the expression in the logarithmic scale. In this context, log ⟨*n*(*E*)⟩ can be interpreted as the entropy for the microcanonical ensemble, as shown in Eq. 9.

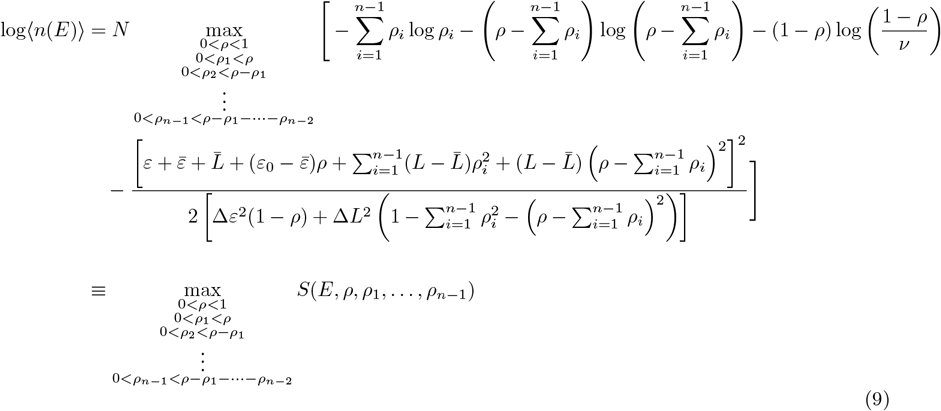

Specifically, 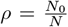denotes the fraction of total native contacts for the entire chain, while 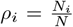 represents the fraction of native contacts in the *i*^th^ native state.

Next, we derive the expressions for energy, entropy, and free energy per residue using the following thermodynamic relation,

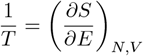

in conjunction with Eq.9. The expression for energy is given by

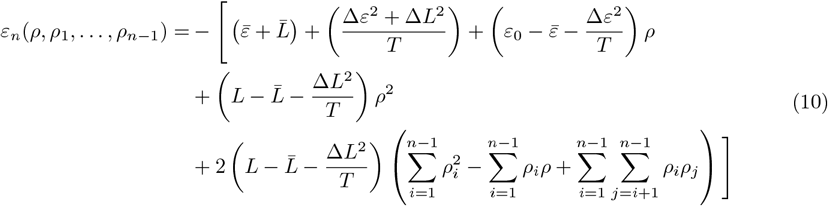

The substitution of the expression from Eq.10 into Eq.9 yields the entropy per residue. The resulting expression is given by:

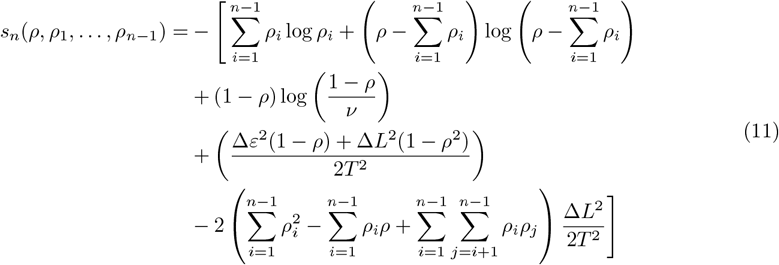

The per-residue Helmholtz free energy *f* is expressed as

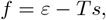

where *ε* and *s* are given by Eq.10 and Eq.11 from our model, respectively. Substituting these expressions into the equation leads to the following expression for *f* :

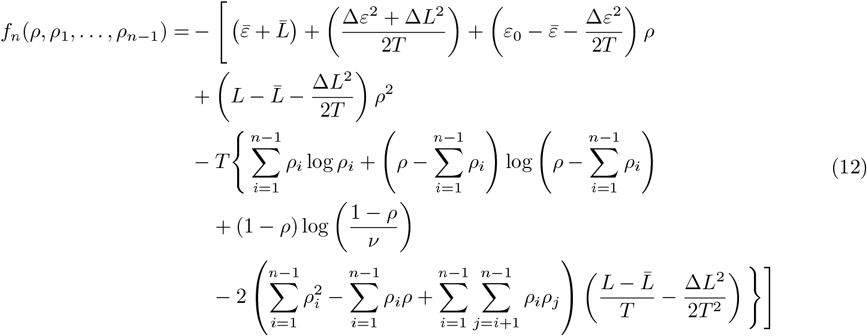

The free energy functional *f* is minimized concerning *ρ* and *ρ*_*i*_ separately to explore the phase diagram and understand the system’s free energy landscape [51, 52]. The necessary partial derivatives are calculated as follows.

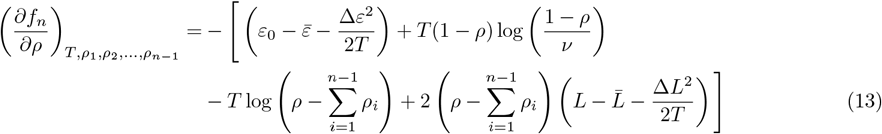

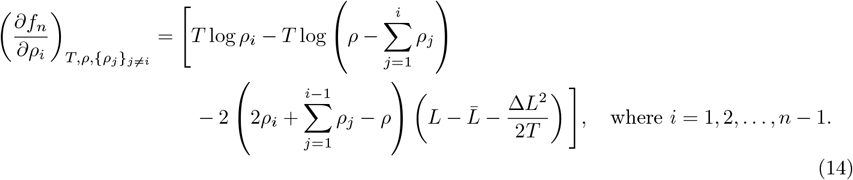

From our model, we also get a critical temperature *T*_0_, where the system’s entropy becomes zero, entering a frozen state, which is also the second-order phase transition temperature for the system.

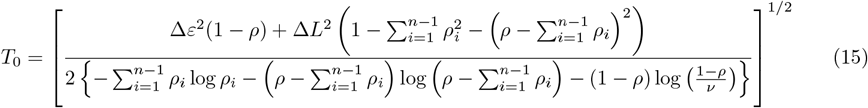

### Biasing a Single Native State over others

In our model, we also examine the effect of stabilizing one of the native states compared to others by introducing a bias factor *δ* into the native energy term at the residue level for one distinct native state. This bias modifies the 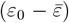 term for residue *N*_1_, thereby stabilizing one native state over others. By incorporating this bias, we can assess its effect on the partial derivatives of the free energy functional *f* with respect to *ρ* and *ρ*_*i*_, as shown in Equations 13 and 14. This analysis is crucial for understanding how stabilizing a single native state influences the order parameter and alters the phase diagram of our multi-native-state system. The modified relevant equations are derived and presented in **S3**.

## Results & Discussion

### Expansion of unfolded region with increasing Native State Multiplicity

The order parameters *ρ* and *ρ*_*i*_ were obtained by solving Equations (13) and (14) simultaneously for systems with native states ranging from *n* = 1 to *n* = 5. In the case of *n* = 1, the model reduces to the single-state scenario (details provided in **S1**), while at *n* = 5, it represents a protein capable of adopting five distinct yet energetically equivalent native structures. We consistently used the parameter values of

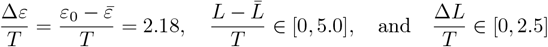

to solve the aforementioned equations. Additionally, we employed a threshold value of *ρ* = 0.75 beyond which the protein can be considered as folded, based on the criteria established by Wolynes and coworkers. [46]. The order parameter *ρ* exhibits two distinct ranges in all systems up to *n* = 5. Values of 0 *< ρ* ≤ 0.2 correspond to the unfolded state of the protein sequence, indicating low levels of native contacts. In contrast, values of 0.75 ≤ *ρ* ≤ 0.95 correspond to the folded state of our model sequence, reflecting the significant formation of native contacts. We constructed a phase diagram by plotting the control parameter 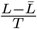 on the *x*-axis and 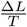 on the *y*-axis. The diagram reveals a clear phase transition line that separates the unfolded and folded regions, effectively distinguishing low and high values of the order parameter *ρ* across all scenarios(Fig. 1(a)). Fig. 1(a) shows a gradual shift in the phase transition line and a contraction of the folded domain as the number of native states increases. The inset highlights *x* ∈ [2.5, 3.25] and *y* ∈ [0, 1.25], where the folded region enclosed by multistate systems is evident. As the complexity rises (i.e., more native states), the area of this folded region in the phase space diminishes, reflecting a lower propensity for the system to remain folded under otherwise identical thermodynamic conditions. In an additional plot where 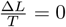, we illustrate how *ρ* varies with the mean deviation of interaction energy, 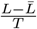 (Fig. 1(b)). This analysis identifies the critical value of 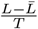 at the onset of the folding transition. As illustrated in Fig. 1(b), the onset points of the phase transition are 2.5, 2.65, 2.8, 2.95, and 3.05 for *n* = 1, 2, 3, 4, and 5, respectively, reflecting a progressive increase consistent with the contraction of the folded domain shown in Fig. 1(a). Notably, at these onset points, the corresponding *ρ* values exceed 0.75, confirming that the overall sequence maintains an overall folded-like fraction of total native contacts.

**Figure 1.**
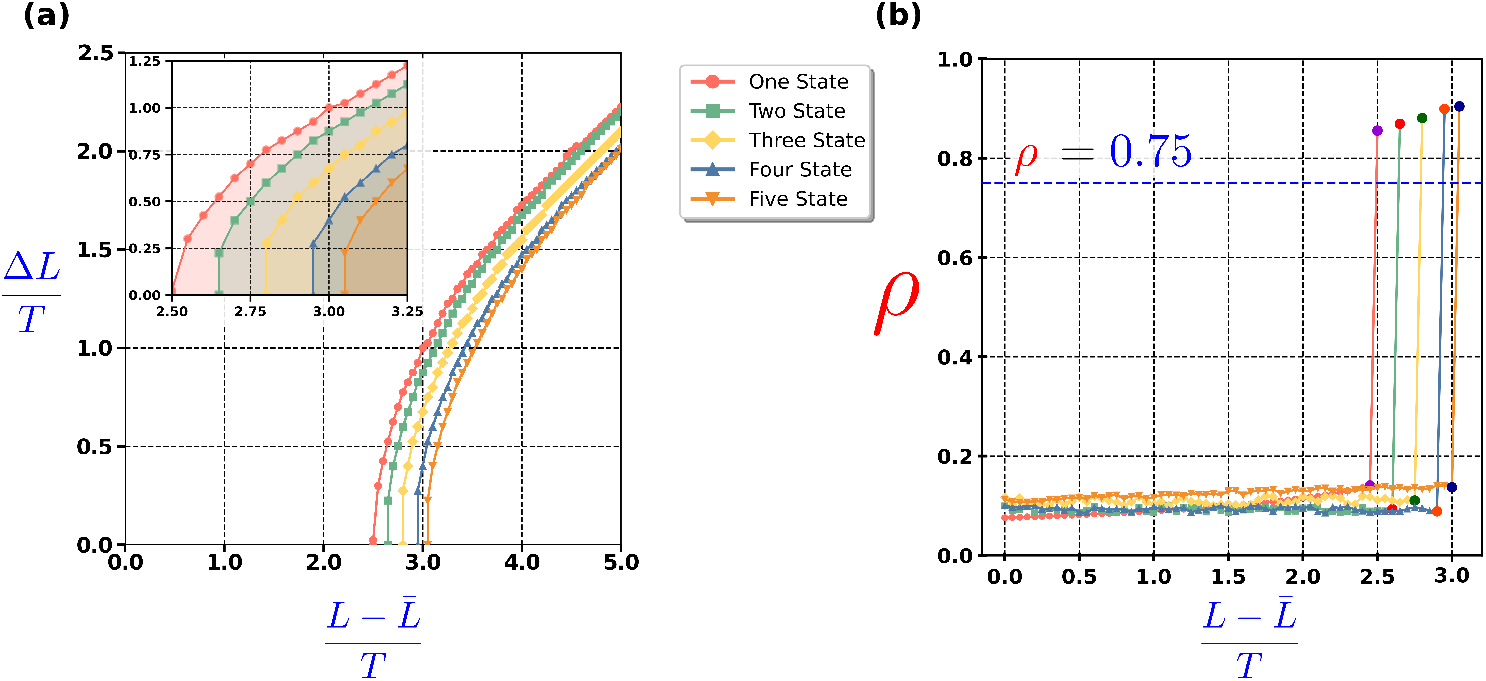
Exploration of protein folding transitions as system complexity increases. **(a)** Phase diagrams for one-to five-state systems, illustrating the contraction of the folded domain as the number of native states *n* increases. The inset highlights the region *x* ∈ [2.5, 3.25] and *y* ∈ [0, 1.25], showcasing the diminishing folded region in multistate systems. **(b)** The variation of the total native contact fraction *ρ* versus 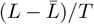 identifies the onset of the phase transition for each *n*. At these onset points, *ρ* for all systems surpass the dotted blue line representing the critical folding threshold *ρ* = 0.75.

### Folding Propensities of individual Native States in Multistate Systems

Next, we determined the individual native contact values *ρ*_*i*_ (with *i* = 1 to 5) while solving Equations (13) and (14), as detailed in the earlier section. Each *ρ*_*i*_ represents the fractional native contacts associated with the *i*-th native state. In the single-state model, the overall fractional native contact *ρ* is identical to *ρ*_1_. However, in systems with multiple states, dividing both sides of Equation (4) by *N* yields the relationship

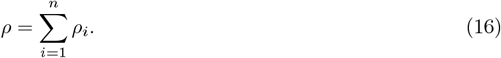

For multi-state systems, the decomposition provides a systematic way to analyze how individual states contribute to the overall folding behavior of the system. We define the highest root as *ρ*_1_ for all systems with varying *n*, followed by the second largest root as *ρ*_2_, and so on for subsequent roots *ρ*_3_, *ρ*_4_, etc. To evaluate the folding behavior of individual native states, we plot *ρ*_1_ against the energy difference 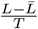 while keeping 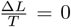. This approach allows us to determine whether the highest *ρ*_1_ value exceeds 0.75, serving as a criterion for identifying folded-like configurations of individual native states relative to the overall folding transition. Additionally, we analyze the variation of other roots against the same parameter to gain further insight into the folding scenario.

Figure 2(a) presents *ρ*_1_ as a function of 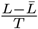 for systems with one to five native states. The plot reveals that for *n* = 1 (single-state), *n* = 2 (two-state), and *n* = 3 (three-state) systems, *ρ*_1_ surpasses the threshold of 0.75 at the transition points along the x-axis. This behavior signifies that at least one of the native states in these systems achieves a folded-like configuration. However, for *n* = 4 and *n* = 5 systems, where the sequence can adopt 4 or 5 equivalent native states, *ρ*_1_ fails to cross the threshold, highlighting a notable change in the folded ensemble. While the overall native contact value *ρ* for all these systems confirms that the system predominantly resides in the folded region of the phase diagram(as shown in Figure 1(b)), the behavior of *ρ*_1_ reveals that for *n* = 4 and *n* = 5, none of the individual native states achieve a folded-like configuration. As shown in **Figure S1**, the *ρ* vs. *ρ*_1_ plot offers additional insight into multi-state folding. The downward trend in the *ρ*–*ρ*_1_ curve for the four- and five-state systems indicates a partial “mixing” scenario, while up to three states exhibit single-state-like folding behavior. This highlights the more intricate folding landscape that emerges when multiple native states are accessible, as contributions to the overall native contacts are distributed among several possible configurations.

**Figure 2.**
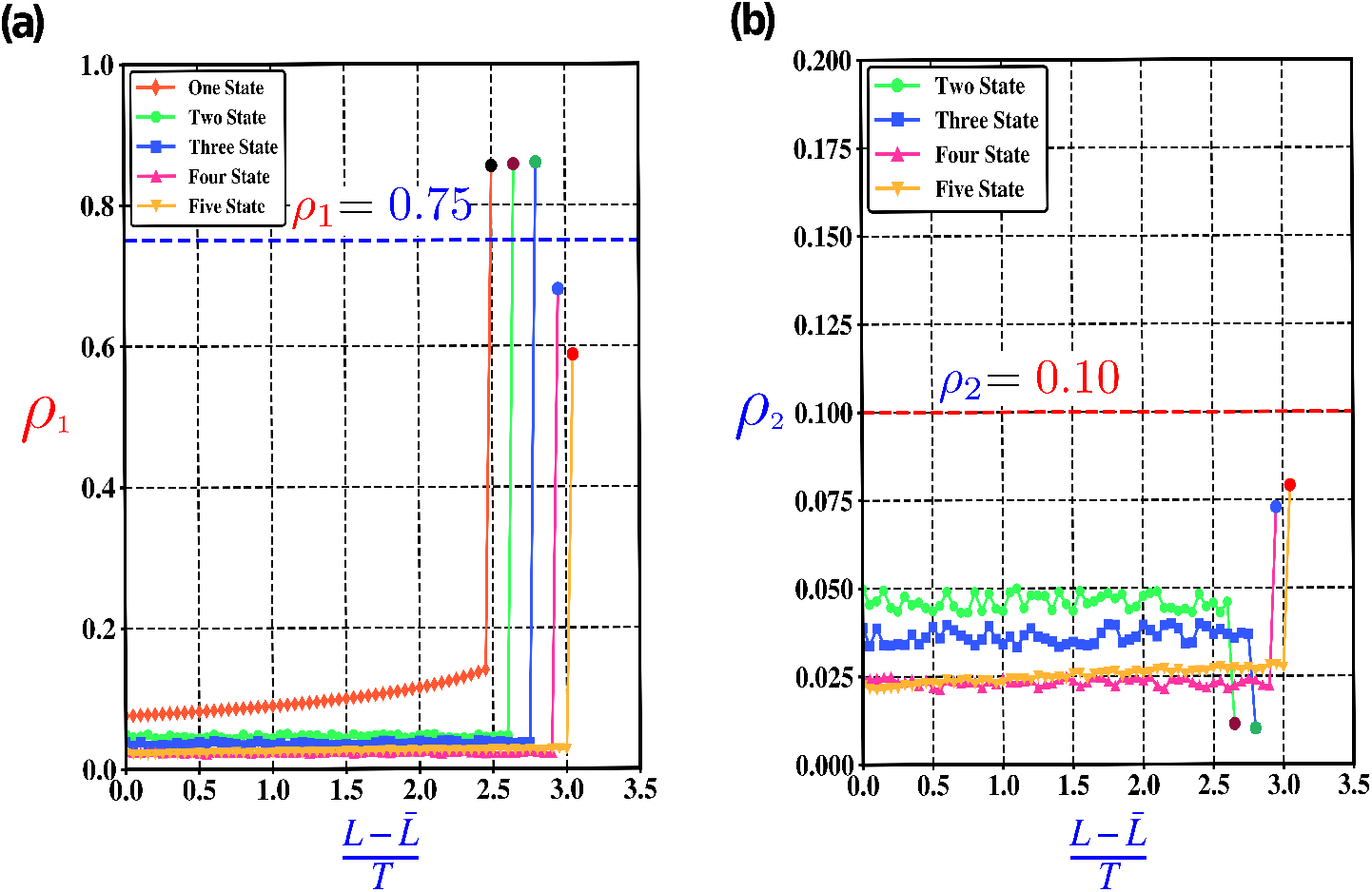
Folding behavior of individual native states. **(a)** Variation of *ρ*_1_, the highest fractional native contact, as a function of 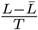. For *n* = 1, 2, and 3, *ρ* crosses the folding threshold indicated by the blue dotted line, while for *n* = 4 and *n* = 5, *ρ*_1_ remains below this line. **(b)** Variation of *ρ*_2_, the second-largest fractional native contact, as a function of 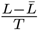. An increasing trend in *ρ* is observed for *n* = 4 and *n* = 5, whereas for *n* = 2 and *n* = 3, *ρ*_2_ remains relatively small, reflecting higher contributions from sub-dominant roots in systems with larger *n*.

Figure 2(b) illustrates the second-largest root, *ρ*_2_, as a function of 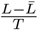 for *n* = 2, 3, 4, and 5 native states. Upon moving from the unfolded to the folded regime, *ρ*_2_ decreases to about 0.01 for *n* = 2 and *n* = 3, indicating a “de-mixing” scenario dominated by a single native state. In contrast, for *n* = 4 and *n* = 5, *ρ*_2_ increases relative to its unfolded value, reaching approximately 0.07 and 0.08, respectively, consistent with a partial “mixing” where multiple states share contributions more uniformly. This trend underscores that, as the number of native states grows, no single conformation achieves a distinctly folded state, leading to a more evenly distributed set of native contacts.

### Folding Landscape for *n* = 2 System

We derived the free energy, entropy, and internal energy expressions for systems with *n* native states, presented in Equations (12), (11), and (10), respectively. For the two native state systems (*n* = 2), the expression of different terms with detailed derivations is provided in **(S2)**. In this case, the system is characterized by two roots, *ρ* and *ρ*_1_, related by *ρ* = *ρ*_1_ + *ρ*_2_. The values of *ρ* and *ρ*_1_ are determined by solving the coupled equations derived by setting *n* = 2 in Equations (13) and (14):

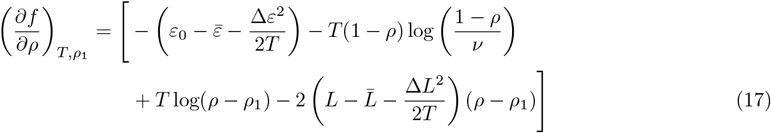

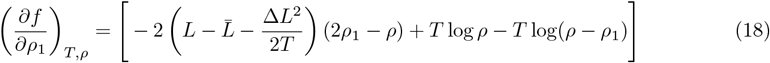

By solving the coupled equations (17) and (18), we obtain a set of values for *ρ* and *ρ*_1_. The folding propensity for the two native-state system, in terms of *ρ* and *ρ*_1_, has been discussed in detail in Figures 1(b) and 2(a), respectively, against the relevant control parameters. For the two-state system, the onset point is found to be 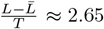 2.65 with 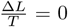, marking the start of the folding transition. We investigate the folding behavior of a two-state system by examining the free energy functional *f*, plotted on the *z*-axis, as a function of the fractional native contacts *ρ* (on the *x*-axis) and *ρ*_1_ (on the *y*-axis), which together characterize the system’s transition between the unfolded and folded states. The analysis is conducted at two distinct values of the control parameter 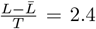 and 2.9, while keeping 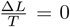. These parameter values are strategically chosen to capture the behavior on either side of the transition point 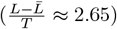: 2.4 in the unfolded regime and 2.9 in the folded regime. Figure 3 showcases the free energy surface of the two native-state systems, capturing its evolution across the unfolded, transition, and folded regimes. Figure 3(a) represents the unfolded regime, where the free energy surface reveals a single stable unfolded state coexisting with two equally stable folded states. Figure 3(b) illustrates the free energy surface at the transition point, where the unfolded and folded states are energetically equivalent. Figure 3(c) corresponds to the folded regime, where the two folded states are energetically more stable than the unfolded state. The free energy surface clearly shows two distinct folded basins: one at (*ρ, ρ*_1_) ≈ (0.868, 0.856) and another at (*ρ, ρ*_1_) ≈ (0.868, 0.012). It follows that for *ρ*_1_ = 0.012, *ρ*_2_ = 0.856. Therefore, the two folded basin are symmetric to each other in terms of order parameters value. For systems with *n* native states, the folded basin will have *n* minima which are symmetric with respect to order parameter values.

**Figure 3.**
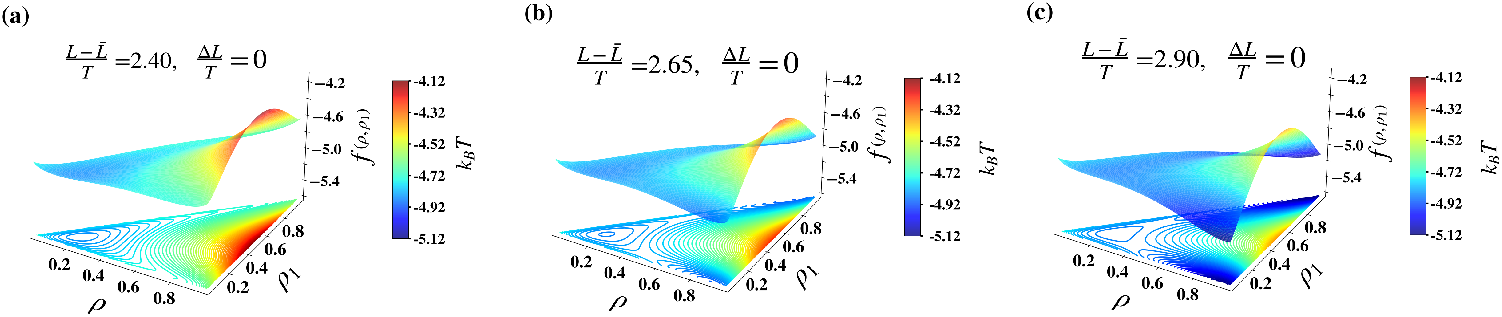
Free energy surfaces depicting the evolution from unfolded to folded regimes. **(a)** A dominant unfolded basin; **(b)** the emergence of two folded basins at the transition point; **(c)** The folded regime is characterized by two equally stable folded basins.

### Modification of the Entropically Frozen Region of the Phase Diagram

In our phase diagram, the second-order phase transition line emerges when the order parameters *ρ, ρ*_*i*_, and Δ*L/T* satisfy the conditions of Equation (11) for an *n*-dimensional system, resulting in zero entropy per residue. Under these conditions, the system enters an entropically frozen state.

To illustrate this behavior, we compare two representative systems: one with a single native folded state and another with two coexisting native folded states. The transition temperature for the one-state system is given by **(Eq.(4) in S1)**, while that for the two-state system is provided by **(Eq.(8) in S2)**.

We systematically compare first-order and second-order phase transitions within a two-dimensional parameter space defined by 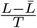and 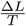. Figure 4 illustrates distinct phase-transition behaviors for systems with one and two native folded states. For *n* = 2, both the folded and frozen regions contract: the red and black lines depict first-order phase boundaries (with the black line corresponding to the two native state system), and its rightward shift indicates contraction of the folded region. The orange boundary, representing the second-order transition line for *n* = 2, shifts leftward—indicating contraction of the frozen region—while the green boundary corresponds to the *n* = 1 system. A shift toward a higher value of 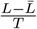 implies a lower transition temperature, and vice versa. Thus, our results indicate a gradual lowering of the folding temperature and an increase in the frozen transition temperature with increasing *n*, consistent with the notion that larger *n* entails greater frustration.

**Figure 4.**
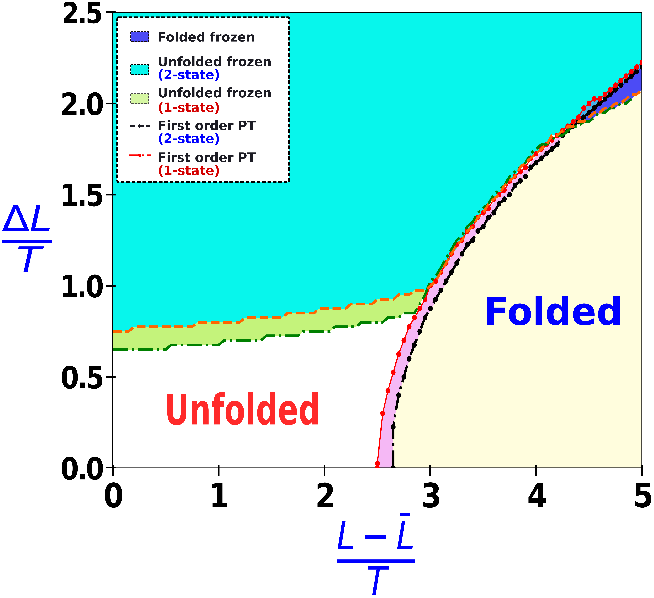
Comparative phase transition analysis for first- and second-order transitions.

### Biasing one native state in n=4 system

We are interested in the four-state system (*n* = 4), where the transitions of *ρ* and *ρ*_1_ have been previously examined in Figures 1(a) and 2(b). It was observed that while *ρ* undergoes a transition into the folded regime at the onset of 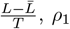 fails to cross the folding threshold of 0.75, indicating that none of the individual native states achieve a folded-like configuration.

To evaluate the impact of biasing one native state out of the possible four, we introduce an additional stability factor *δ* to the residue level interaction energy term *ε − ε*_0_ of one native state. This controlled bias alters the stability of the selected native state relative to others, as reflected in the modified free energy functional (**Eq.(9) in S3**). We determine the modified roots, *ρ* and *ρ*_*i*_, by solving the three coupled equations (**Equations (10), (11), and (12) in S3**),with the parameter *δ* explicitly included in the corresponding equation. This approach allows us to assess how the perturbation affects the stability and native contact values of individual states.

Figure 5(a) displays the phase diagram for both the biased and unbiased (*n* = 4) systems at *δ ≈* 0.4. The perturbation shifts the phase transition line to the left, resulting in an expanded folding region compared to unbiased one. The larger folding region in the perturbed system suggests an increased folding propensity. Additionally, Figure 5(b) shows that the perturbed *ρ*_1_ exceeds the critical threshold of 0.75, while the unperturbed *ρ*_1_ remains below this value. This indicates that perturbations enhance the folding propensity of selected individual native state and folds into that particular native state.

**Figure 5.**
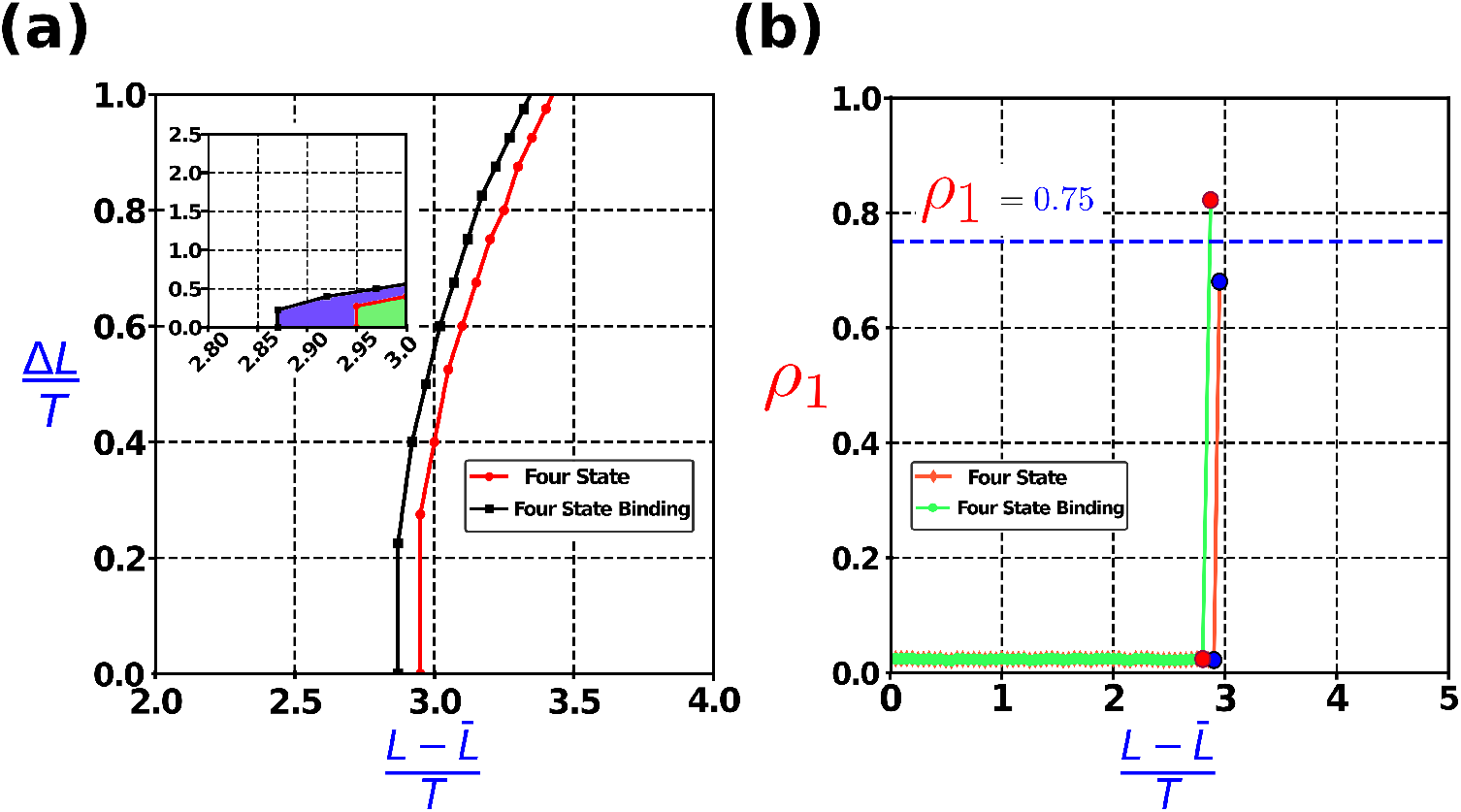
**(a)** Phase diagrams for four-state systems under biased and unbiased conditions. The main panel shows that bias shifts the phase transition line leftward and enlarges the folding region. The inset compares the biased state (black line with blue folded region) to the unbiased state (red line with green folded region), with the blue region being notably larger. **(b)** *ρ*_1_ transitions show that only the perturbed system’s *ρ*_1_ crosses the blue dotted folding cutoff line, indicating enhanced folding propensity.

## Discussion and Conclusion

Our study demonstrates that increasing the number of native folded states in protein models leads to a contraction of the folded domain, indicating a decrease in the folding propensity of the system. Additionally, there are two scenarios that exist. In the first scenario, for *n* = 1 to *n* = 3, where both overall native fraction *ρ* and the highest individual native fraction, *ρ*_1_, surpass the folding threshold of 0.75. More specifically, there exist two folded minima in the *n* = 2 system and three folded minima in the *n* = 3 system where a fraction of native contacts of one distinct native state crosses the threshold value. This scenario can be related to the proteins having more than one folded state as in the metamorphic proteins.

In the second scenario, for *n* = 4 and *n* = 5, the expansion of the unfolded region becomes quite significant. This suggests that such sequences with larger values of *n* will find themselves in the unfolded region of the phase diagram under physiological thermodynamic conditions. This scenario parallels with the intrinsically disorder proteins where the configurations remain unfolded-like in physiological conditions. Additionally, in the folded configuration of such sequences, none of the individual native states achieve a distinctly folded configuration.

To examine the effect of selective stabilization, we introduce a targeted stabilization of one particular native state in the *n* = 4 system. This modification, results in expansion of the folded region in the phase diagram compared to the unperturbed case, increasing the folding propensity of the sequence. Additionally, the fractional native contact value of the stabilized native state exceeds the folding threshold, a scenario absent in the unperturbed system, indicating a folded configuration similar to *n* = 1 case. This result demonstrates how selective stabilization can enhance folding propensity at the level of individual native states, offering a potential understanding of the folding upon binding scenario of the intrinsically disorder proteins.

Our study provides a theoretical framework for understanding folding scenarios in protein system with multiple possible native states, emphasizing how native-state multiplicity shapes folding phase diagram, metamorphic scenario and the coupled folding upon binding scenario of the IDPs.

## Conflicts of Interest

The authors declare that they have no conflict of interest with respect to the publication of this article.

## Acknowledgments

The authors sincerely thank the Central Supercomputing Facility of the Indian Association for the Cultivation of Science, Kolkata and DST (DST-SERB grant **CRG/2020/000756**) for providing essential computational resources. Rohon Mitra gratefully acknowledges the Council for Scientific and Industrial Research (CSIR) for the award of a research fellowship, which supported this work.

## Notes

### Competing Interest Statement

The authors have declared no competing interest.

## References

[1] Christian B. Anfinsen. Principles that govern the folding of protein chains. Science, 181(4096):223– 230, 1973.

[2] E. E. Lattman and G. D. Rose. Protein folding–what’s the question? Proceedings of the National Academy of Sciences, 90(2):439–441, 1993.

[3] Seiji Saito and Masaki Sasai. Kinetic consistency in the protein folding processdedicated to professor keiji morokuma in celebration of his 65th birthday.1. Journal of Molecular Structure: THEOCHEM, 461-462:503–521, 1999.

[4] Andrej Šali, Eugene Shakhnovich, and Martin Karplus. How does a protein fold? Nature, 369:248–251, 1994.

[5] Rohit V. Pappu, Rajgopal Srinivasan, and George D. Rose. The flory isolated-pair hypothesis is not valid for polypeptide chains: Implications for protein folding. Proceedings of the National Academy of Sciences, 97(23):12565–12570, 2000.

[6] José Nelson Onuchic and Peter G Wolynes. Theory of protein folding. Current Opinion in Structural Biology, 14(1):70–75, 2004.

[7] D. Thirumalai, Edward P. Brien, Greg Morrison, and Changbong Hyeon. Theoretical perspectives on protein folding. Annual Review of Biophysics, 39(Volume 39, 2010):159–183, 2010.

[8] David Balchin, Manajit Hayer-Hartl, and F. Ulrich Hartl. In vivo aspects of protein folding and quality control. Science, 353(6294):aac4354, 2016.

[9] R Zwanzig, A Szabo, and B Bagchi. Levinthal’s paradox. Proceedings of the National Academy of Sciences, 89(1):20–22, 1992.

[10] Ken A Dill and Hue Sun Chan. From levinthal to pathways to funnels. Nature Structural Biology, 4(1):10–19, 1997.

[11] Peter G. Wolynes. Folding funnels and energy landscapes of larger proteins within the capillarity approximation. Proceedings of the National Academy of Sciences, 94(12):6170–6175, 1997.

[12] Hugh Nymeyer, Angel E. García, and José Nelson Onuchic. Folding funnels and frustration in off-lattice minimalist protein landscapes. Proceedings of the National Academy of Sciences, 95(11):5921–5928, 1998.

[13] H. Frauenfelder, S. G. Sligar, and P. G. Wolynes. The energy landscapes and motions of proteins. Science, 254(5038):1598–1603, 1991.

[14] J. D. Bryngelson, J. N. Onuchic, N. D. Socci, and P. G. Wolynes. Funnels, pathways, and the energy landscape of protein folding: A synthesis. Proteins: Structure, Function, and Bioinformatics, 21(3):167–195, 1995.

[15] José Nelson Onuchic, Zaida Luthey-Schulten, and Peter G. Wolynes. Theory of protein folding: The energy landscape perspective. Annual Review of Physical Chemistry, 48(Volume 48, 1997):545–600, 1997.

[16] Hue Sun Chan and Ken A. Dill. Protein folding in the landscape perspective: Chevron plots and non-arrhenius kinetics. Proteins: Structure, Function, and Bioinformatics, 30(1):2–33, 1998.

[17] Saikat Dhibar and Biman Jana. Optimized collective variable for collapse transition in linear hydrophobic polymers: Importance of hydration water and end-to-end distance. Journal of Chemical Theory and Computation, 20(17):7404–7415, 2024. PMID: 39252562.

[18] Nicholas C. Fitzkee and George D. Rose. Steric restrictions in protein folding: An α-helix cannot be followed by a contiguous β-strand. Protein Science, 13(3):633–639, 2004.

[19] John Jumper, Richard Evans, Alexander Pritzel, Tim Green, Michael Figurnov, Olaf Ronneberger, Kathryn Tunyasuvunakool, Russ Bates, Augustin Žídek, Anna Potapenko, Alex Bridgland, Clemens Meyer, Simon A. A. Kohl, Andrew J. Ballard, Andrew Cowie, Bernardino Romera-Paredes, Stanislav Nikolov, Rishub Jain, Jonas Adler, Trevor Back, Stig Petersen, David Reiman, Ellen Clancy, Michal Zielinski, Martin Steinegger, Michalina Pacholska, Tamas Berghammer, Sebastian Bodenstein, David Silver, Oriol Vinyals, Andrew W. Senior, Koray Kavukcuoglu, Pushmeet Kohli, and Demis Hassabis. Highly accurate protein structure prediction with alphafold. Nature, 596(7873):583–589, 2021.

[20] Ken A. Dill and Justin L. MacCallum. The protein-folding problem, 50 years on. Science, 338(6110):1042–1046, 2012.

[21] William A. Eaton and Peter G. Wolynes. Theory, simulations, and experiments show that proteins fold by multiple pathways. Proceedings of the National Academy of Sciences, 114(46):E9759– E9760, 2017.

[22] S. Walter Englander and Leland Mayne. The nature of protein folding pathways. Proceedings of the National Academy of Sciences, 111(45):15873–15880, 2014.

[23] Christopher M Dobson. Protein folding and misfolding. Nature, 426(6968):884–890, 2003.

[24] Pengfei Tian and Robert B. Best. Structural determinants of misfolding in multidomain proteins. PLOS Computational Biology, 12(5):1–28, 05 2016.

[25] Shaik Basha, Darshan Chikkanayakanahalli Mukunda, Jackson Rodrigues, Meagan Gail D’Souza, Gireesh Gangadharan, Aparna Ramakrishna Pai, and Krishna Kishore Mahato. A comprehensive review of protein misfolding disorders, underlying mechanism, clinical diagnosis, and therapeutic strategies. Ageing Research Reviews, 90:102017, 2023.

[26] Weihua Zheng, Min-Yeh Tsai, and Peter G. Wolynes. Comparing the aggregation free energy landscapes of amyloid beta(1–42) and amyloid beta(1–40). Journal of the American Chemical Society, 139(46):16666–16676, 2017. PMID: 29057654.

[27] Mandira Dutta, Michael R. Diehl, José N. Onuchic, and Biman Jana. Structural consequences of hereditary spastic paraplegia disease-related mutations in kinesin. Proceedings of the National Academy of Sciences, 115(46):E10822–E10829, 2018.

[28] Debasis Saha and Biman Jana. Identifying the template for oligomer to fibril conversion for amyloid-β (1-42) oligomers using hamiltonian replica exchange molecular dynamics. ChemPhysChem, 23(24):e202200393, 2022.

[29] Dennis J. Selkoe. Cell biology of protein misfolding: The example of alzheimer’s and parkinson’s diseases. Nature Cell Biology, 6(11):1054–1061, 2004.

[30] F. Ulrich Hartl. Protein misfolding diseases. Annual Review of Biochemistry, 86(Volume 86, 2017):21–26, 2017.

[31] Itamar Yadid, Noam Kirshenbaum, Michal Sharon, Orly Dym, and Dan S. Tawfik. Metamorphic proteins mediate evolutionary transitions of structure. Proceedings of the National Academy of Sciences, 107(16):7287–7292, 2010.

[32] Acacia F. Dishman, Robert C. Tyler, Jamie C. Fox, Andrew B. Kleist, Kenneth E. Prehoda, M. Madan Babu, Francis C. Peterson, and Brian F. Volkman. Evolution of fold switching in a metamorphic protein. Science, 371(6524):86–90, 2021.

[33] Catherine Ghosh and Biman Jana. Curious case of mad2 protein: Diverse folding intermediates leading to alternate native states. The Journal of Physical Chemistry B, 126(9):1904–1916, 2022. PMID: 35230837.

[34] Sergey V. Solomatin, Max Greenfeld, Steven Chu, and Daniel Herschlag. Multiple native states reveal persistent ruggedness of an rna folding landscape. Nature, 463(7281):681–684, 2010.

[35] Jeffrey K. Noel, Alexander Schug, Abhinav Verma, Wolfgang Wenzel, Angel E. Garcia, and José N. Onuchic. Mirror images as naturally competing conformations in protein folding. The Journal of Physical Chemistry B, 116(23):6880–6888, 2012. PMID: 22497217.

[36] Joseph M. Rogers, Annette Steward, and Jane Clarke. Folding and binding of an intrinsically disordered protein: Fast, but not ’diffusion-limited’. Journal of the American Chemical Society, 135(4):1415–1422, 2013. PMID: 23301700.

[37] Jiaan Yang, Wen-xiang Cheng, Gang Wu, Sitong Sheng, and Peng Zhang. Prediction of folding patterns for intrinsic disordered protein. Scientific Reports, 13(1):20343, 2023.

[38] Srinivasaraghavan Kannan, David P. Lane, and Chandra S. Verma. Long range recognition and selection in idps: the interactions of the c-terminus of p53. Scientific Reports, 6(1):23750, 2016.

[39] Amit Kumar, Prateek Kumar, Shobha Kumari, Vladimir N. Uversky, and Rajanish Giri. Folding and structural polymorphism of p53 c-terminal domain: One peptide with many conformations. Archives of Biochemistry and Biophysics, 684:108342, 2020.

[40] S. F. Edwards and P. W. Anderson. Theory of spin glasses. Journal of Physics F: Metal Physics, 5(5):965–974, May 1975.

[41] David Sherrington and Scott Kirkpatrick. Solvable model of a spin-glass. Phys. Rev. Lett., 35:1792–1796, Dec 1975.

[42] G. Parisi. The order parameter for spin glasses: A function on the interval 0-1. Journal of Physics A: Mathematical and General, 13(3):1101–1112, Mar 1980.

[43] M Mezard, G Parisi, and M Virasoro. Spin Glass Theory and Beyond. WORLD SCIENTIFIC, 1986.

[44] K. Binder and A. P. Young. Spin glasses: Experimental facts, theoretical concepts, and open questions. Rev. Mod. Phys., 58:801–976, Oct 1986.

[45] Patrick Charbonneau, Enzo Marinari, Marc Mézard, Giorgio Parisi, Federico Ricci-Tersenghi, Gabriele Sicuro, and Francesco Zamponi. Spin Glass Theory and Far Beyond. WORLD SCIENTIFIC, 2023.

[46] J. D. Bryngelson and P. G. Wolynes. Spin glasses and the statistical mechanics of protein folding. Proceedings of the National Academy of Sciences, 84(21):7524–7528, 1987.

[47] L. Olivares-Quiroz and L. S. Garcia-Colin. Protein’s unfolding and the glass transition: a common thermodynamic signature. AIP Conference Proceedings, 978(1):109–114, 02 2008.

[48] Thomas C. T. Michaels, Daoyuan Qian, and Andela Šaric. Amyloid formation as a protein phase transition. Nature Reviews Physics, 5(7):379–397, 2023.

[49] Biman Jana, Dwaipayan Chakrabarti, and Biman Bagchi. Glassy orientational dynamics of rodlike molecules near the isotropic-nematic transition. Phys. Rev. E, 76:011712, Jul 2007.

[50] R. A. Goldstein, Z. A. Luthey-Schulten, and P. G. Wolynes. Optimal protein-folding codes from spin-glass theory. Proceedings of the National Academy of Sciences, 89(11):4918–4922, 1992.

[51] Erik M. Boczko and Charles L. Brooks. First-principles calculation of the folding free energy of a three-helix bundle protein. Science, 269(5222):393–396, 1995.

[52] Pradipta Kumar Das and Biman Jana. Registry alteration in dynein’s microtubule-binding domain: A aaa domain-guided event. Chemical Physics Impact, 9:100702, 2024.

