## Supplementary Information for "A model of protein folding with multiple native states: Metamorphicity, Intrinsic disorderness and folding upon binding of proteins"

### 1 Supplementary Equations

#### S1: Thermodynamic Equations for the Single-State ($n = 1$) Model

$$\varepsilon = \left[ -(\bar{\varepsilon} + \bar{L}) - \left(\frac{\Delta\varepsilon^2 + \Delta L^2}{T}\right) - \left(\varepsilon_0 - \bar{\varepsilon} - \frac{\Delta\varepsilon^2}{T}\right)\rho - \left(L - \bar{L} - \frac{\Delta L^2}{T}\right)\rho^2 \right] \tag{1}$$

$$s = \left[ -\rho \log\rho - (1-\rho)\log\!\left(\frac{1-\rho}{\nu}\right) - \left(\frac{\Delta\varepsilon^2\,(1-\rho) + \Delta L^2\,(1-\rho^2)}{2\,T^2}\right) \right] \tag{2}$$

$$\begin{aligned} f = \Bigg[ & -\bar{\varepsilon} - \bar{L} - \left(\frac{\Delta\varepsilon^2 + \Delta L^2}{2\,T}\right) - \left(\varepsilon_0 - \bar{\varepsilon} - \frac{\Delta\varepsilon^2}{2\,T}\right)\rho - \left(L - \bar{L} - \frac{\Delta L^2}{2\,T}\right)\rho^2 \\ & + T\,\rho\log\rho + T\,(1-\rho)\log\!\left(\frac{1-\rho}{\nu}\right) \Bigg] \end{aligned} \tag{3}$$

$$T_0 = \left[ \frac{\Delta\varepsilon^2 + \Delta L^2 - \Delta\varepsilon^2\,\rho - \Delta L^2\,\rho^2}{2\left\{ -\rho\log\rho - (1-\rho)\log\!\left(\frac{1-\rho}{\nu}\right)\right\}} \right]^{1/2} \tag{4}$$

The derived equations, obtained by setting $n = 1$ in our model with multiple native states, yield expressions for internal energy, entropy, free energy, and transition temperature. Specifically, Eqs. (1), (2), (3), and (4) describe these thermodynamic quantities in the single-native-state limit. These expressions are consistent with those presented by Wolynes and coworkers in their seminal work [1] , demonstrating the agreement of our framework with established theoretical results.

#### S2: Two-State ($n = 2$) Analogue and Thermodynamic Framework

$$\varepsilon(\rho, \rho_1) = -\Bigg[ \left(\bar{\varepsilon} + \bar{L}\right) + \left(\frac{\Delta\varepsilon^2 + \Delta L^2}{T}\right) + \left(\varepsilon_0 - \bar{\varepsilon} - \frac{\Delta\varepsilon^2}{T}\right)\rho + \left(L - \bar{L} - \frac{\Delta L^2}{T}\right)\left(\rho_1^2 + (\rho - \rho_1)^2\right)\Bigg] \tag{5}$$

$$s(\rho, \rho_1) = \Bigg[ -\rho_1 \log \rho_1 - (\rho - \rho_1)\log(\rho - \rho_1) - (1-\rho)\log\left(\frac{1-\rho}{\nu}\right) - \left\{\frac{\Delta\varepsilon^2(1-\rho) + \Delta L^2(1 - \rho_1^2 - (\rho - \rho_1)^2)}{2T^2}\right\}\Bigg] \tag{6}$$

$$f(\rho, \rho_1) = \Bigg[ -\bar{\varepsilon} - \bar{L} - \left(\frac{\Delta\varepsilon^2 + \Delta L^2}{2T}\right) - \left(\varepsilon_0 - \bar{\varepsilon} - \frac{\Delta\varepsilon^2}{2T}\right)\rho - \left(L - \bar{L} - \frac{\Delta L^2}{2T}\right)\rho_1^2 - \left(L - \bar{L} - \frac{\Delta L^2}{2T}\right)(\rho - \rho_1)^2 + T\rho_1 \log \rho_1 + T(\rho - \rho_1)\log(\rho - \rho_1) + T(1-\rho)\log\left(\frac{1-\rho}{\nu}\right)\Bigg] \tag{7}$$

$$T_0 = \left[\frac{\Delta\varepsilon^2(1-\rho) + \Delta L^2\left(1 - \rho_1^2 - (\rho - \rho_1)^2\right)}{2\left\{-\rho_1 \log \rho_1 - (\rho - \rho_1)\log(\rho - \rho_1) - (1-\rho)\log\left(\frac{1-\rho}{\nu}\right)\right\}}\right]^{1/2} \tag{8}$$

#### S3: Single-State Bias ($\delta$), Modified Free Energy and Partial Derivatives

$$f_n(\rho, \rho_1, \ldots, \rho_{n-1}) = -\Bigg[ \left(\bar{\varepsilon} + \bar{L}\right) + \left(\frac{\Delta\varepsilon^2 + \Delta L^2}{2T}\right) + \left\{(\varepsilon_0 - \bar{\varepsilon})(1+\delta)\rho_1 + (\varepsilon_0 - \bar{\varepsilon})(\rho - \rho_1) - \frac{\Delta\varepsilon^2}{2T}\rho\right\} + \left(L - \bar{L} - \frac{\Delta L^2}{2T}\right)\rho^2 - T\Bigg\{\sum_{i=1}^{n-1}\rho_i \log \rho_i + \left(\rho - \sum_{i=1}^{n-1}\rho_i\right)\log\left(\rho - \sum_{i=1}^{n-1}\rho_i\right) + (1-\rho)\log\left(\frac{1-\rho}{\nu}\right) - 2\left(\sum_{i=1}^{n-1}\rho_i^2 - \sum_{i=1}^{n-1}\rho\rho_i + \sum_{i=1}^{n-1}\sum_{j=i+1}^{n-1}\rho_i\rho_j\right)\left(\frac{L - \bar{L}}{T} - \frac{\Delta L^2}{2T^2}\right)\Bigg\}\Bigg] \tag{9}$$

$$
\left(\frac{\partial f_n}{\partial \rho}\right)_{T,\rho_1,\rho_2,\ldots,\rho_{n-1}} = - \left[ \left(\varepsilon_0 - \bar{\varepsilon} - \frac{\Delta\varepsilon^2}{2T}\right) + T(1-\rho)\log\left(\frac{1-\rho}{\nu}\right) \right.
$$

$$
\left. - \; T\log\left(\rho - \sum_{i=1}^{n-1}\rho_i\right) + 2\left(\rho - \sum_{i=1}^{n-1}\rho_i\right)\left(L - \bar{L} - \frac{\Delta L^2}{2T}\right)\right], \tag{10}
$$

$$
\left(\frac{\partial f_n}{\partial \rho_1}\right)_{T,\rho,\rho_2,\rho_3,\ldots,\rho_{n-1}} = \left[ -(\varepsilon_0 - \bar{\varepsilon})\delta + T\log\rho_1 - T\log\left(\rho - \rho_1\right) \right.
$$

$$
\left. -(2\rho_1 - \rho) \; 2\left(L - \bar{L} - \frac{\Delta L^2}{2T}\right)\right] \tag{11}
$$

$$
\left(\frac{\partial f_n}{\partial \rho_i}\right)_{T,\rho,\{\rho_j\}_{j\neq i}} = \left[ T\log\rho_i - T\log\left(\rho - \sum_{j=2}^{i}\rho_j\right) \right.
$$

$$
\left. -\left(2\rho_i + \sum_{j=2}^{i-1}\rho_j - \rho\right) 2\left(L - \bar{L} - \frac{\Delta L^2}{2T}\right)\right], \quad \text{where } i = 2, 3, \ldots, n-1. \tag{12}
$$

### 2 Supplementary Figures

#### $\rho$ vs. $\rho_1$: Folded-State Propensity in Multi-State Models

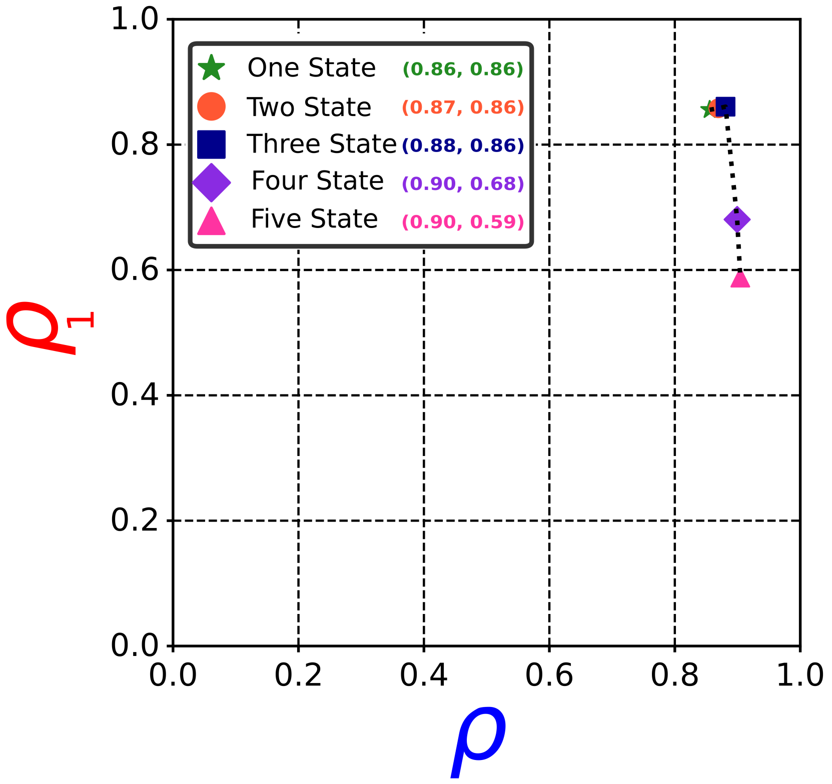

**Figure S1:** Plot of $\rho$ vs. $\rho_1$ across systems with multiple native folded states, illustrating reduced folding propensity in four- and five-state models where no native state surpasses the 0.75 threshold for classification as folded.

### References

[1] J. D. Bryngelson and P. G. Wolynes. Spin glasses and the statistical mechanics of protein folding. *Proceedings of the National Academy of Sciences*, 84(21):7524–7528, 1987.
